# Multimodal CustOmics: A Unified and Interpretable Multi-Task Deep Learning Framework for Multimodal Integrative Data Analysis in Oncology

**DOI:** 10.1101/2024.01.20.576363

**Authors:** Hakim Benkirane, Maria Vakalopoulou, David Planchard, Julien Adam, Ken Olaussen, Stefan Michiels, Paul-Henry Cournède

## Abstract

Characterizing cancer poses a delicate challenge as it involves deciphering complex biological interactions within the tumor’s microenvironment. Histology images and molecular profiling of tumors are often available in clinical trials and can be leveraged to understand these interactions. However, despite recent advances in representing multimodal data for weakly supervised tasks in the medical domain, numerous challenges persist in achieving a coherent and interpretable fusion of whole slide images and multi-omics data. Each modality operates at distinct biological levels, introducing substantial correlations both between and within data sources. In response to these challenges, we propose a deep-learning-based approach designed to represent multimodal data for precision medicine in a readily interpretable manner. While demonstrating superior performance compared to state-of-the-art methods across multiple test cases, our approach also provides robust results and extracts various scores characterizing the activity of each modality and their interactions at the pathway and gene levels. The strength of our method lies in its capacity to unravel pathway activation through multimodal relationships and extend enrichment analysis to spatial data for supervised tasks. We showcase the efficiency and robustness of its predictive capacity and interpretation scores through an extensive exploration of multiple TCGA datasets and validation cohorts, underscoring its value in advancing our understanding of cancer. The method is publicly available in [Github](https://github.com/HakimBenkirane/Multimodal-CustOmics).

## 1 Introduction

Deep learning in medical tasks has gradually gained acceptance in the scientific community^1–3^. In many cases, cancer diagnosis, prognosis, and therapeutic response prediction rely on heterogeneous data to comprehensively characterize the patient’s profile^4–6^. To acquire the maximum amount of information for performing those tasks, it is often necessary to integrate data sources corresponding to different levels of biological processes, like histology images, molecular profiles, and clinical data. However, even though deep learning has revolutionized computer vision in medical imaging^7,8^, the integration of multiple sources of data is still an open challenge. Because of the high dimensionality of histology images and multi-omics data, learning an interpretable multimodal representation that can leverage the interactions between the different sources of data and adapt to downstream predictive tasks is complex.

Numerous studies and projects have made available datasets from various molecular sources. Integrating different omics data can reduce uncertainty arising from varying experimental conditions and unveil interactions beyond the reach of a single source. The inclusion of multiple molecular sources raises critical dimensionality issues due to the abundance of genes or biomarkers. It also comes with substantial heterogeneity due to the intricate genomic complexity of the human molecular profile. Furthermore, each molecular source may encapsulate distinct biological functions that hold critical importance for extracting clinically relevant insights from the resulting analyses.

In computational pathology, gigapixel Whole-Slide images (WSIs) have become the golden standard for prediction tasks^9^. However, their enormous resolution makes supervised tasks dependent on computationally heavy training and appropriate representations. Most works on using gigapixel images for weakly supervised tasks consist of Multiple Instance Learning (MIL) formulations^10,11^. Small patches from the WSI are extracted as independent instances and converted into instance-level representations before being pooled using a global aggregation over the bag of unordered instances. However, this data representation does not capture the spatial context, making modeling heterogeneous visual concepts in the tumor micro-environment difficult.

In this work, we will present a new way of integrating histology images with the patient’s molecular profile in a representation that captures high correlations between and within different dimensions of the biological system (i.e., biological pathways, cellular regions…). This framework uses a multiple-instance variational autoencoder with modality dropout to integrate multimodal data while remaining robust to missing data. One of the key contributions is to adapt the autoencoder to create a mixture of expert representation similar to Jordan et al.^12^ between multiple relevant biological dimensions of the patient’s profile that will enhance interpretability. This yields three significant advantages:

- The division of each modality by different functional dimensions, we achieve a general understanding of a needle-in-a-haystack problem where all modalities are highly dimensional.
- By implementing modality dropout within the framework of our method, we can create an end-to-end pipeline that is robust to missing modalities.
- Because of the multidimensional form of the representation, interactions between all the dimensions from the different modalities are easily interpretable thanks to the mixture of expert networks.

## 2 Results

### 2.1 Representation Learning of Multimodal Data

To address the different challenges of integrating multimodal data, we propose a deep-learning framework to create an interpretable image-omic representation that captures interactions at multiple levels of the biological system. Called CustOmics, this integration method relies on the strategy introduced in Benkirane et al.^13^. The original network optimally integrated heterogeneous data from different omics sources while conserving each modality’s individuality. The new version of our multimodal network can now jointly integrate H&E slides and molecular profile features (mutation status, copy-number variation, RNA sequencing [RNA-seq] expression, DNA methylation…). Moreover, it can also interpret how the interaction between all those sources correlates with specific supervised tasks like molecular subtype identification or survival prediction. Moreover, this method allows the assessment of feature importance at multiple levels through ad-hoc scores. At the gene level, the method outputs the importance score of each gene for each molecular source, both independently and in association with other modalities. At the pathway level, a Multimodal Pathway Enrichment Score (MPES) is computed to assess the importance of a specific pathway for a specific prediction task, such as molecular subtype classification or survival outcomes. This score has also been extended to account for spatial correlations through the histopathology slide and reflects the importance of the interaction between spatial regions of the WSI and pathways.

We compare CustOmics to four other methods integrating omics and histopathology data. For this comparison, we follow the study conducted by Chen et al.^14^ and use as a basis of comparison the engineered baselines introduced for omics, histopathology, and multi-omics integration:

#### SNN

As a multi-omics baseline, we train a feed-forward self-normalizing network architecture^15^ where multi-omics data are concatenated before being fed to the network. This architecture is used in^14,16^ and is state-of-the-art for histology-genomic integration.

#### DeepSets

One of the first neural architectures for set-based deep learning problems^17^, which propose sum pooling over instance level features. Its multimodal extension is presented in^14^, where the omics data are processed with an SNN and integrated with bilinear pooling.

#### Attention MIL

A set-based network similar to DeepSets replaces the sum pooling by an attention pooling technique^18^.

#### DeepAttnMISL

A set-based network that first applies K-Means clustering to instance-level features, processing each cluster using Siamese networks and then aggregating the cluster features using global attention pooling.^19^.

#### MCAT

The current state-of-the-art in multimodal histology-genomics integration, presented in^14^. It is a transformer-based set-based network that combines modalities with bilinear attention-based pooling.

In this study, we implemented a multi-level interpretability approach, focusing on gene and spatial levels. Spatial interpretability results were further explained by extracting high-attention image patches and analyzing them with the pre-trained Hover-Net model for cell instance segmentation and classification^20^. This analysis classified cells into tumors, lymphocytes, stromal, necrosis, and epithelial cells. We quantitatively evaluated the frequency of these cell types in high-attention patches for each patient. Additionally, we estimated the proportion of Tumor Infiltrating Lymphocytes (TILs) in these patches using the methodology developed by Saltz et al.^21^, providing a deeper understanding of the tumor microenvironment.

### 2.2 Multimodal Datasets for Multi-Task Predictions

In this study, we use the pan-cancer dataset extracted from the Genomic Data Commons (GDC)^22^ with 11,768 patients and 33 tumor types and containing multi-omics data, histopathology slides, and clinical data. We evaluate the performances of our model both on the whole pan-cancer dataset and on smaller cohorts of specific tumor types to show the robustness of our method to a smaller number of patients. The goal is to evaluate CustOmics on both the classification of tumor types and the prediction of survival outcomes. For the survival prediction task, eight cohorts were selected. This selection was made according to the number of patients and the censoring rate. Details of those cohorts are presented in Table S1. Among those eight cohorts, three were also used to evaluate the classification into molecular subtypes: TCGA-BRCA, TCGA-COAD, and TCGA-STAD. To validate the interpretability results provided by our method, we use a validation cohort based on the International Adjuvant Lung Cancer Trial (IALT)^23^, regrouping patients with non-small-cell lung cancer with histopathology slides, mutation, and copy-number variations data (More details on the validation set can be found in Table S2). The validation is done on a survival outcome prediction using histopathology slides and DNAseq data. We compare the results of this validation dataset to the TCGA-LUAD and TCGA-LUSC cohorts.

### 2.3 CustOmics provides improved predictions in multiple predictive tasks

Multiple test cases were executed in a comprehensive evaluation of CustOmics within a multitask framework encompassing both classification and survival analyses. The performances across survival and classification tasks are presented in Tables 1 and 2.

**Table 1.** Classification Perfomances: Comparison of the classification performances for the 4 tasks with respect to the area under ROC (AUC %).

| Methods | PANCAN | BRCA | COAD | STAD |
| --- | --- | --- | --- | --- |
| SNN (Multi-Omics) | 94.1 ± 2.7 | 92.0 ± 3.3 | 79.2 ± 3.3 | 84.6 ± 3.3 |
| CustOmics (Multi-Omics) | **98.9 ± 1.4** | **98.3 ± 1.0** | **88.1 ± 1.9** | **98.4 ± 1.2** |
| DeepSets (WSI Only) | 84.7 ± 3.3 | 68.6 ± 4.1 | 55.2 ± 4.3 | 58.9 ± 4.6 |
| AttnMIL (WSI Only) | 88.4 ± 2.0 | 71.2 ± 4.9 | 56.2 ± 4.4 | 61.4 ± 4.4 |
| DeepAttnMISL (WSI Only) | 89.8 ± 2.5 | 71.1 ± 3.3 | 55.7 ± 4.0 | 62.1 ± 4.5 |
| MCAT (WSI Only) | 90.4 ± 1.8 | 72.3 ± 3.3 | 61.7 ± 3.3 | 69.4 ± 3.5 |
| CustOmics (WSI Only) | **92.5 ± 1.2** | **73.2 ± 3.1** | **62.2 ± 2.0** | **71.4 ± 2.2** |
| DeepSets (WSI + Multi-Omics) | 96.7 ± 1.5 | 84.9 ± 2.0 | 58.6 ± 2.2 | 67.1 ± 2.7 |
| AttnMIL (WSI + Multi-Omics) | 97.1 ± 1.2 | 86.6 ± 2.1 | 60.0 ± 2.5 | 69.4 ± 2.7 |
| DeepAttnMISL (WSI + Multi-Omics) | 97.8 ± 1.1 | 88.3 ± 2.7 | 65.4 ± 2.7 | 66.6 ± 2.2 |
| MCAT (WSI + Multi-Omics) | 98.9 ± 1.1 | 95.4 ± 2.0 | 93.3 ± 1.2 | 88.7 ± 2.3 |
| CustOmics (WSI + Multi-Omics) | **99.5 ± 0.9** | **98.7 ± 1.1** | **94.7 ± 1.0** | **96.3 ± 2.4** |

**Table 2.** Survival Perfomances: Comparison of the survival performances for the 8 TCGA cohorts with respect to the Concordance Index (C-index %).

| Methods | BLCA | BRCA | COAD | GBMLGG | KIRC | LUAD | STAD | UCEC |
| --- | --- | --- | --- | --- | --- | --- | --- | --- |
| SNN (Multi-Omics) | 54.6 ± 2.7 | 47.5 ± 4.9 | 50.8 ± 3.4 | 60.5 ± 3.1 | 59.0 ± 5.7 | 54.2 ± 5.4 | 51.1 ± 4.6 | 49.6 ± 8.3 |
| **CustOmics (Multi-Omics)** | **64.1 ± 1.5** | **63.4 ± 1.8** | **58.6 ± 1.8** | **78.4 ± 1.6** | **66.5 ± 2.3** | **63.7 ± 1.2** | **55.6 ± 1.4** | **68.5 ± 2.8** |
| DeepSets (WSI Only) | 50.6 ± 5.1 | 50.1 ± 6.5 | 50.2 ± 3.9 | 50.9 ± 3.5 | 49.6 ± 5.4 | 50.7 ± 7.5 | 50.0 ± 7.5 | 50.3 ± 7.5 |
| AttnMIL (WSI Only) | 54.8 ± 4.8 | 57.0 ± 5.7 | 59.6 ± 3.9 | 79.0 ± 3.2 | 56.9 ± 6.1 | 56.3 ± 6.8 | 57.8 ± 6.1 | 63. ± 7. |
| DeepAttnMISL (WSI Only) | 50.3 ± 5.5 | 52.1 ± 5.9 | 53.1 ± 3.3 | 73.8 ± 3.9 | 56.9 ± 6.6 | 55.0 ± 6.1 | 58.8 ± 5.3 | 60.0 ± 7.2 |
| MCAT (WSI Only) | 55.5 ± 3.2 | 57.1 ± 5.6 | **59.9 ± 2.5** | **79.4 ± 2.0** | 56.0 ± 3.4 | 55.0 ± 6.1 | 57.7 ± 3.9 | 63.4 ± 6.7 |
| **CustOmics (WSI Only)** | **56.7 ± 3.9** | **59.4 ± 3.7** | 58.5 ± 2.1 | 78.7 ± 2.0 | **61.3 ± 2.2** | **60.0 ± 5.5** | **59.6 ± 2.3** | **67.9 ± 2.2** |
| DeepSets (WSI + Multi-Omics) | 59.6 ± 4.7 | 52.1 ± 7.1 | 61.4 ± 3.6 | 81.8 ± 3.3 | 54.2 ± 5.4 | 56.8 ± 7.3 | 52.9 ± 5.8 | 59.0 ± 7.4 |
| AttnMIL (WSI + Multi-Omics) | 57.4 ± 5.3 | 54.8 ± 6.4 | 61.9 ± 3.5 | 81.3 ± 3.0 | 62.0 ± 6.2 | 58.5 ± 6.7 | 54.0 ± 6.2 | 56.9 ± 6.1 |
| DeepAttnMISL (WSI + Multi-Omics) | 58.5 ± 5.4 | 58.0 ± 7.6 | 61.0 ± 3.2 | 81.4 ± 3.3 | 60.7 ± 7.1 | 55.0 ± 6.1 | 53.8 ± 5.7 | 59.1 ± 6.6 |
| MCAT (WSI + Multi-Omics) | 62.4 ± 3.6 | 58.3 ± 5.5 | 62.8 ± 2.9 | 82.5 ± 2.3 | 66.5 ± 3.9 | 62.5 ± 4.5 | 56.2 ± 3.1 | 62.2 ± 2.7 |
| **CustOmics (WSI + Multi-Omics)** | **67.2 ± 2.5** | **65.2 ± 3.6** | **64.4 ± 2.0** | **84.2 ± 2.3** | **68.2 ± 2.1** | **64.9 ± 3.7** | **58.0 ± 1.5** | **68.0 ± 2.2** |

The initial assessment involved cancer-type classification within the TCGA Pancancer cohort, revealing CustOmics’ superior performance in terms of AUC compared to other benchmarked methods^24^. This notable performance can be attributed to the substantial patient cohort size and the rich information embedded within molecular data, a phenomenon well-documented across multiple studies^13,25,26^.

To underscore the method’s robustness concerning sample size, evaluations were conducted on smaller datasets, focusing on predicting molecular subtypes within three specific TCGA cohorts: TCGA-BRCA, TCGA-COAD, and TCGA-STAD. Across all classification tasks (as detailed in Table 1), CustOmics consistently outperformed other comparative methods. Notably, in multi-omics integration, the mixed-integration VAE within CustOmics demonstrated superior performance compared to the SNN utilized by other methods. An additional assessment showcased the significance of multi-omics integration, delineated in Table S4, emphasizing the impact on performance when replacing the VAE encoder in CustOmics with a standard SNN.

Further exploration into the integration strategies revealed the advantage of CustOmics in exploiting diverse molecular data sources. Comparative analyses in Table S3 demonstrated that while SNN performed best with RNAseq alone, CustOmics exhibited enhanced performance when integrating RNAseq with CNV and methylation data. This divergence in integration strategies suggests CustOmics’ capacity to augment predictive power and unveil novel interactions among disparate data sources.

The evaluation extended to survival outcome prediction across eight TCGA cohorts, consistently showcasing CustOmics’ superior predictive capabilities compared to alternative methods.

Notably, in survival analysis based solely on Whole Slide Images (WSI), CustOmics exhibited comparatively weaker performance in specific cohorts compared to the transformer architecture employed in MCAT. An ablation study (detailed in Table S4) replaced CustOmics’ hypergraph encoding for WSI embeddings with a visual transformer, yielding superior performance with the hypergraph embeddings for multimodal representation learning.

In survival tasks, particularly in multi-omics scenarios (without WSI), CustOmics displayed substantially more significant differences in performances, especially as other methods like SNN showed concordance indices approaching randomness (Table 2). CustOmics’ lower standard deviation across folds underscores its enhanced robustness compared to other state-of-the-art approaches.

### 2.4 CustOmics can evaluate pathway enrichment for the discrimination of Her2 subtype in breast cancer

CustOmics conducts pathway enrichment analysis across multiple tasks to comprehend its output comprehensively. Figure 2 presents interpretability findings concerning the PAM50 subtype classification within the TCGA-BRCA dataset. The objective is to elucidate the determinants driving the discrimination of specific subtypes, notably the Her2 subtype, within a multimodal context.

**Figure 1.**
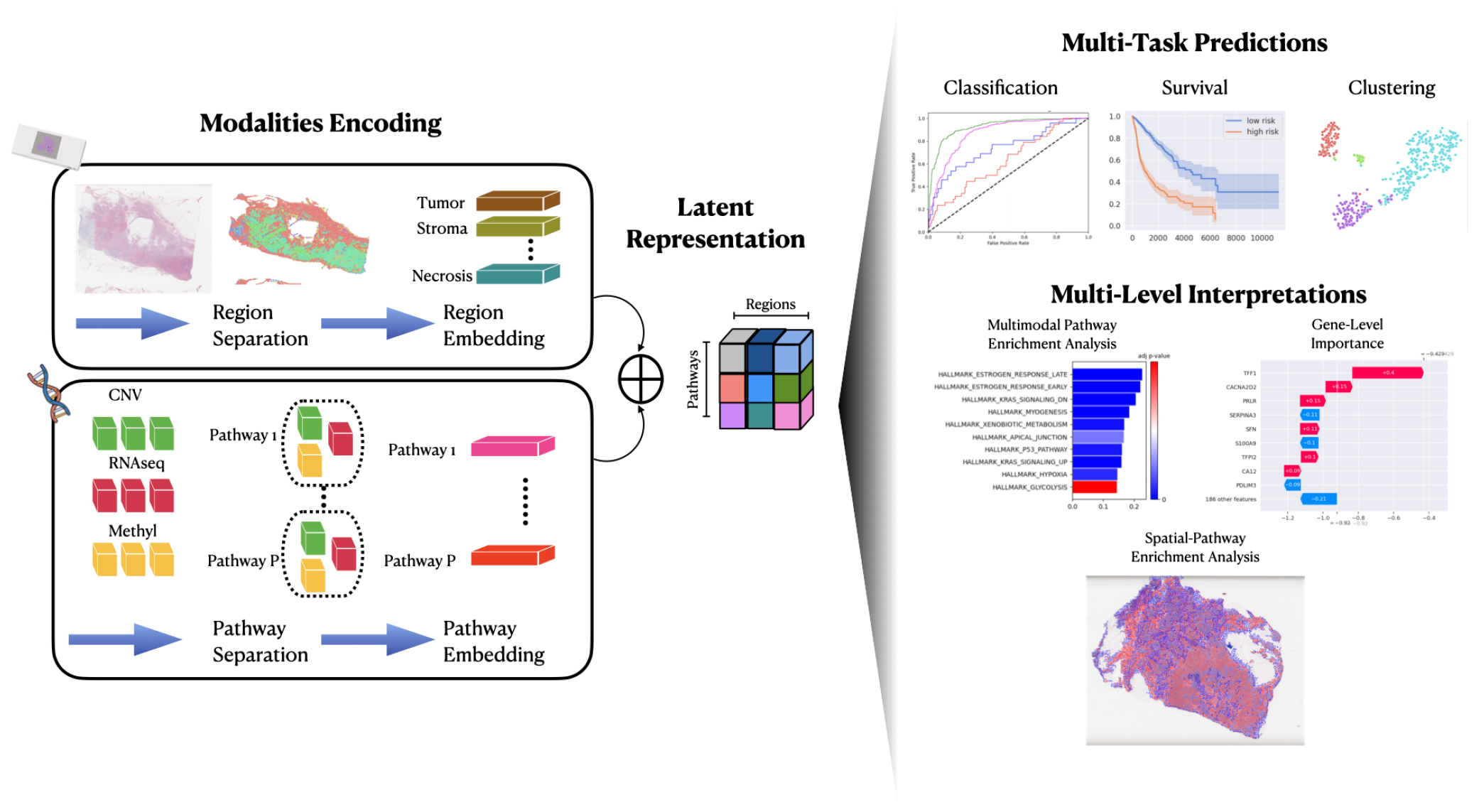
**a. Modalities Encoding:** Each modality is encoded using a specific methodology. For histopathology slides, spatial regions are extracted using a hypergraph encoder so that we can obtain an embedding for each region. For multi-omic integration, genes are separated by pathways, which gives place to a multi-omics embedding per pathway. **b.Latent Representation:** The latent representation is comprised of multiple blocks, each block being the embedding of the interaction between a region and a pathway. **c. Multimodal Task:** The latent representation is then used either for supervised tasks like classification or survival analysis, or in an unsupervised fashion for clustering for example. **c. Multi-Level Interpretations:** Interpretation Results are extracted at multiple levels: Gene-level, pathway-level and spatial-level

**Figure 2.**
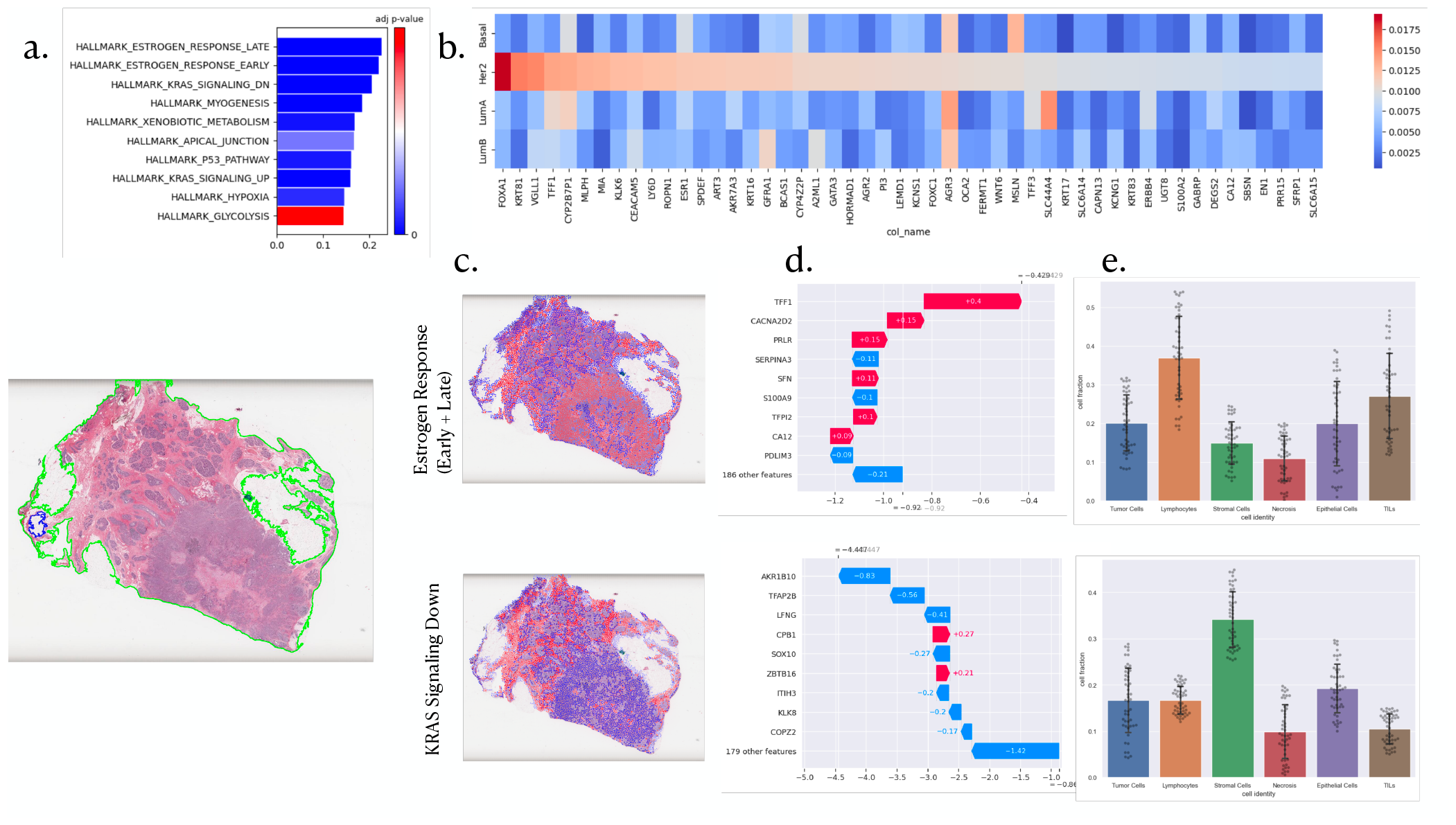
PAM50 interpretability analysis: **a**. Pathway enrichment scores and p-value associated with the GSVA. **b**. SHAP values for the most important genes influencing the stratification of the Her2 subtype and their influence on other subtypes. **c**. Spatial Enrichment Analysis for the top 2 pathways and their most important genes. **d**. Gene importance inside the considered pathways. **e**. Cell distribution in the top 10% attention regions.

The initial layer of interpretability operates at the gene level, utilizing a Multi-Omics Pathway Enrichment Score (MPES) and conducting Gene Set Variation analysis using normalized gene importance scores (detailed in the methods section). Figure 2b delineates essential pathways in Her2 subtype discrimination, notably highlighting the significance of estrogen response and KRAS signaling down hallmarks. The interrelation between the Her2 subtype and estrogen response has been extensively investigated^27^, emphasizing their coexpression’s multifaceted impact on breast carcinogenesis, invasive behavior, and cellular growth.

Further exploration into gene-level importance is depicted in Figure 2c, spotlighting the predominant genes responsible for discriminating the Her2 subtype and contrasting their importance across other subtypes. Notably, the FOXA1 gene emerges with substantial importance, aligning with its suggested role as a transcription factor for Her2, as indicated in Cruz et al.^28^.

Beyond multi-omics pathway enrichment analysis, CustOmics extends interpretability to encompass multimodal enrichment, revealing spatial interactions within histopathology slides that correlate with specific pathways for discriminating Her2 subtypes. Figure 2d showcases such interpretability outcomes for the estrogen response and KRAS pathways. A description of different cell populations within high-importance regions for each pathway is provided to enhance comprehension. Notably, regions associated with the KRAS pathway exhibit heightened proportions of stromal cells, suggesting potential regulation of tumor cell signaling via these stromal cells^29^. Conversely, the estrogen response demonstrates a strong interaction with regions featuring elevated densities of Tumor-Infiltrating Lymphocytes (TILs) and lymphocytes, corroborated by multiple sources^30,31^. This interaction holds particular significance when stratifying between ER- and ER+ patients.

### 2.5 CustOmics can evaluate pathway enrichment for Pancancer Survival Analysis

In a similar vein, interpretability analysis extends to survival analysis. Figure 3 delineates the varying degrees of enrichment analysis for predicting survival outcomes within the TCGA Pancancer dataset.

**Figure 3.**
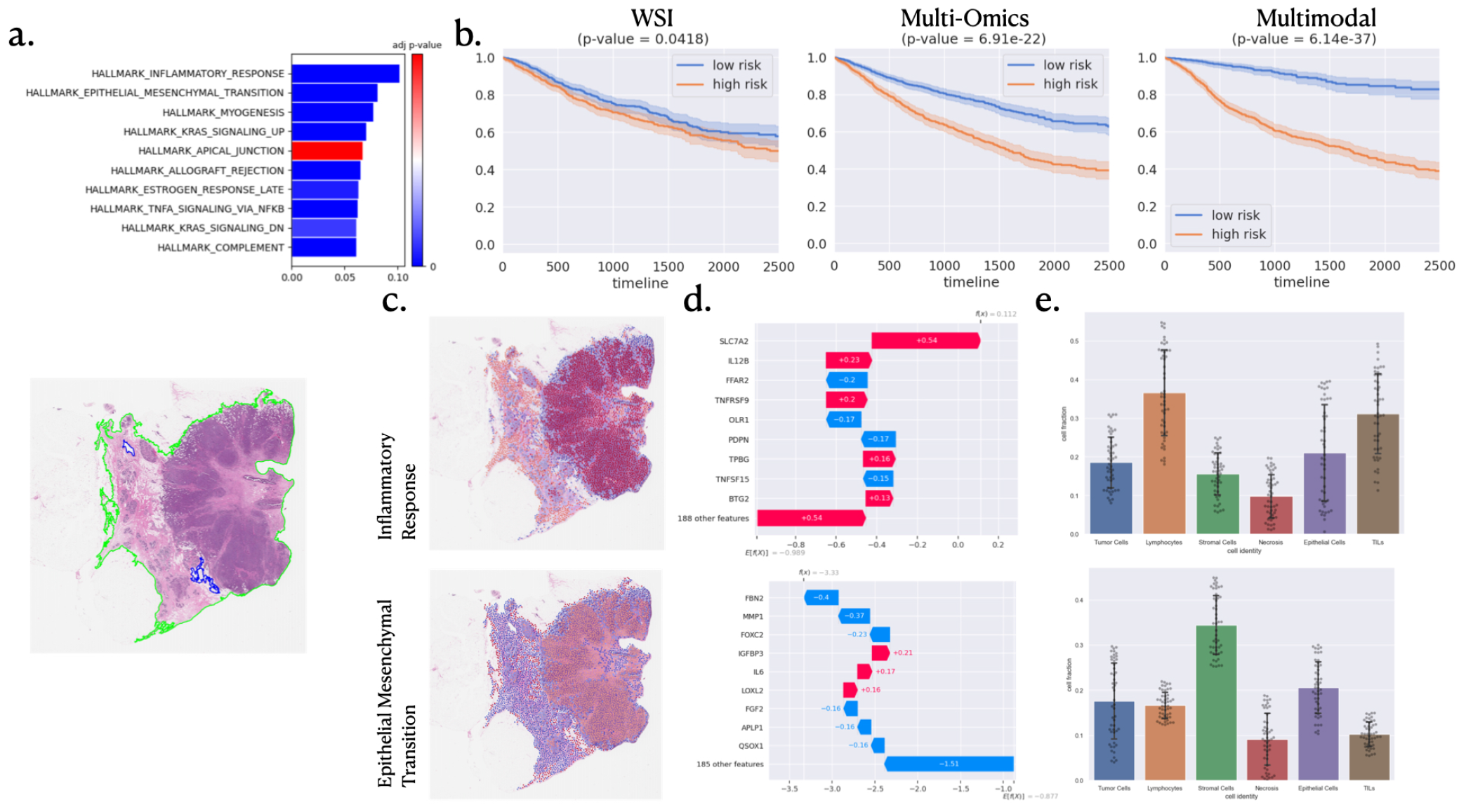
**a**. Pathway enrichment analysis for the pan-cancer survival outcome prediction task. **b**. Kaplan Meier curve associated with the pan-cancer survival outcome prediction task with a computed log-rank p-value for the stratification of high and low risk for death event. **c**. Spatial Enrichment Analysis for the top 2 pathways and their most important genes. **d**. Gene importance inside the considered pathways. **e**. Cell distribution in the top 10% attention regions.

Specifically, Figure 3b underscores the predominant influence of the inflammatory response pathway on survival outputs, a finding consistent with existing literature^32^.

Demonstrating the relevance of employing multimodal integration in pan-cancer survival analysis, Figure 3c illustrates the impact of incorporating multiple data sources on stratifying low and high-risk patients, as evidenced by Kaplan-Meier curves and their corresponding log-rank p-values.

Further investigation into the effect of essential pathways is portrayed in Figure 3d, showcasing the significance of interactions between the two most crucial pathways. Notably, heightened importance is observed within the inflammatory response pathway, characterized by increased lymphocyte densities and Tumor-Infiltrating Lymphocytes (TILs). In contrast, the epithelial-mesenchymal transition pathway manifests greater densities of stromal cells.

Delving deeper into the Epithelial-Mesenchymal Transition pathway, the primary influential gene appears to be FBN2, renowned for its inhibition of cancer cell invasion and migration, as stated in Mahdizadehi et al.^33^, thereby explaining its inclination toward lower risk outcomes.

### 2.6 CustOmics shows robust interpretability results for Lung Cancer

In order to ascertain the reliability of interpretability findings by Customics, validation was conducted employing a dataset from the IALT study focusing on lung cancer. A comparative analysis of survival analysis interpretability results between the IALT dataset and a combined dataset of TCGA-LUAD/TCGA-LUSC was undertaken.

The pathway enrichment analysis, illustrated in Figure 4, revealed congruent outcomes between the TCGA and IALT datasets, identifying KRAS signaling down as the foremost pathway. This finding aligns with established literature^34^, which underscores the prevalence of oncogenic KRAS mutations in approximately 25% of lung adenocarcinoma cases, thus representing a pivotal focus in current drug development.

**Figure 4.**
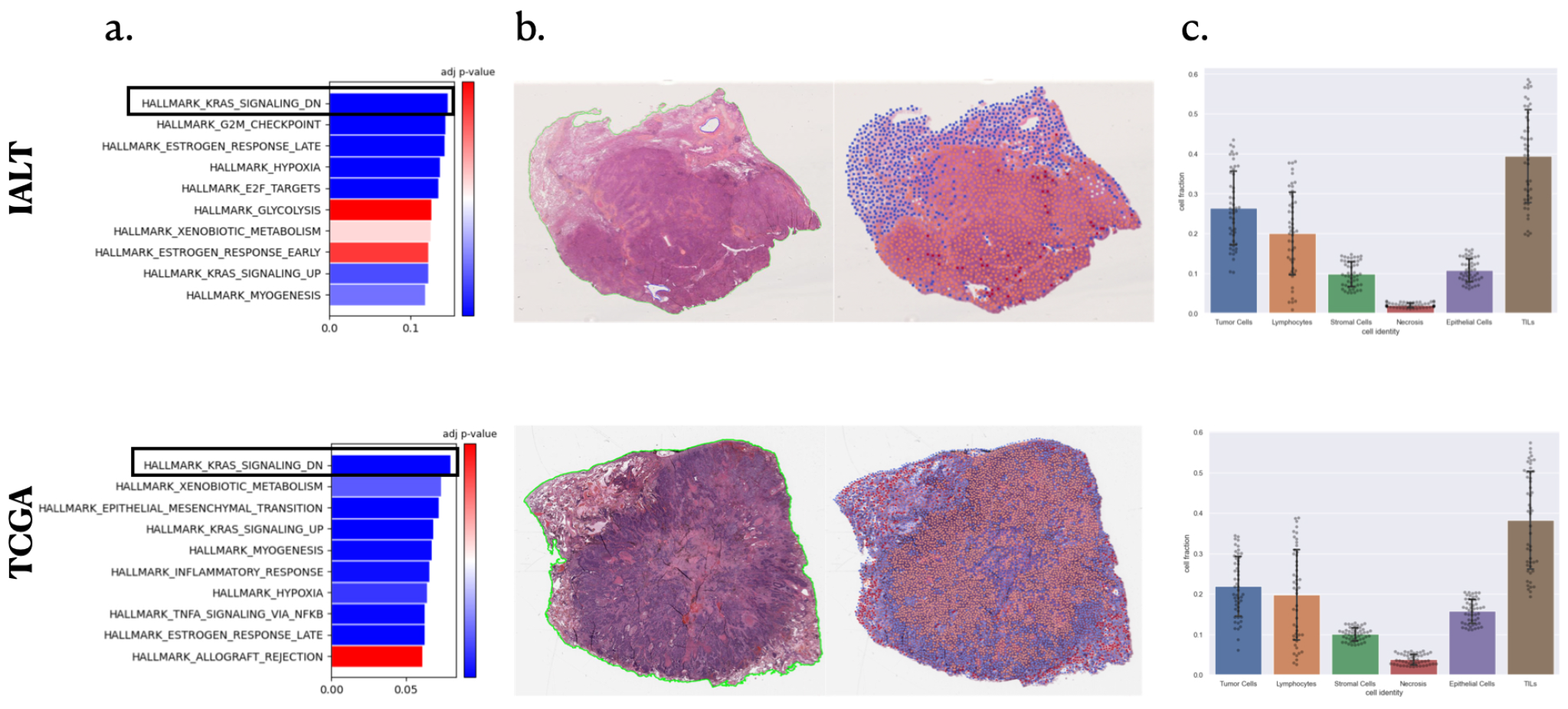
Comparison of interpretations between the TCGA-LUAD/TCGA-LUSC datasets and IALT regarding the distinction between high and low survival risk. **a**. Pathway Enrichment analysis highlighting top pathways for the task. **b**. Spatial Importance of the KRAS Signaling Down pathway, illustrating regions of high importance in interaction with this pathway. **c**. Comparison of cell populations within high importance regions of the Whole Slide Image (WSI).

Exploration of the interplay between the KRAS signaling down pathway and spatial regions within histopathology images was subsequently pursued. Figure 4b depicts that regions of high attention in both datasets exhibit similar distributions of cell types, indicating the robustness of our method across distinct datasets of the same cancer type. Notably, this distribution underscores the association between the KRAS pathway and tumor-infiltrating lymphocytes (TILs) heightened densities and marginally increased tumoral cell counts in predicting survival outcomes. This consistent association echoes previous findings in^35^, highlighting a solid correlation between KRAS mutation status and tumor immunity-related characteristics, notably CD8+ TILs.

## 3 Discussion

The CustOmics framework represents a comprehensive toolset to bridge prediction and interpretation within biological systems across multiple levels: genes, pathways, and spatial orientations. This multifaceted system generates three distinct interpretability scores concurrent with predictions, unraveling the biological knowledge underlying model outcomes. Empirical assessments underscore CustOmics’ robust predictive capabilities, outperforming state-of-the-art methodologies in integrating multi-omics and histopathology data across eight diverse datasets. However, despite its efficiency with smaller datasets in this study, CustOmics’ reliance on deep learning methodologies might restrict efficiency when confronted with limited training data availability.

Furthermore, CustOmics is distinguished by its interpretability, facilitating a broader spectrum of analyses, thereby enriching comprehension across diverse biological modalities. Notably, although this study centered on three omics data types, CustOmics exhibits versatility in seamlessly integrating varied omics data without necessitating framework alterations. This adaptability originates from an initial phase that independently trains each source, serving as a normalization layer for heterogeneous sources.

The strategic segmentation of inputs into interpretable entities for both spatial and molecular data mitigates challenges arising from the high dimensionality of Whole Slide Images (WSIs) and multi-omics datasets. This partitioning augments interpretability and broadens the method’s applicability to uncharted pathways or spatial regions beyond this study’s scope.

CustOmics places significant emphasis on expansive interpretability functionalities. This aims to unveil predominant biological functions steering specific predictions across diverse data sources and scales. This comprehensive approach fosters collaboration between biologists and computational pathologists, offering an enriched framework for in-depth analyses through enrichment analysis for omics and spatial data. The method extracts cohesive insights from diverse data sources, unveiling a panoramic view of interconnected biological processes influencing predictive outcomes. The integrated interpretation strategy serves as a common ground for interdisciplinary collaboration, empowering researchers to explore the complexities of biological systems effectively. By integrating omics and spatial data within enrichment analysis, CustOmics enables a deeper understanding of the interplay between molecular information and spatial contexts, enriching investigative pathways for researchers in the field.

Indeed, in the scope of this research, several limitations are noteworthy. Firstly, while this study successfully delineates interactions between omics and histopathology data, a method to effectively discern and capitalize on the individual contributions of each omic source to a patient’s molecular profile remains absent. This absence inhibits a comprehensive understanding of each omic source’s distinct impact on the overall molecular landscape. Secondly, the link established between the generated representation and phenotype data, beyond mere predictive labels, solely relies on the conditioning of the latent space. While this conditioning methodology effectively incorporates phenotypical signals into the multimodal representation, it lacks a mechanism to explicitly unveil interaction insights between different modalities and diverse clinical variables. This omission presents an avenue for future development, potentially enhancing the interpretability of multimodal interactions and their associations with clinical factors.

## 4 Methods

Within the scope of this study, we design, implement and evaluate a multimodal integration network for integrating histopathology slides and multi-omics data. For 1≤ *i* ≤ *N*, let us denote by *W*_*i*_ and *O*_*i*_ respectively the WSI and multi-omics bag for patient *i*. The goal of this study is to build and train a multi-task network *ℳ* that takes as input the two bags and create an interpretable multimodal representation *z*_*i*_ for each patient such that *ℳ* (*W*_*i*_, *O*_*i*_) = *z*_*i*_.

### 4.1 Representation of Histopathology Images

Initially, an automated tissue segmentation is conducted using the method outlined in^36^, which uses Otsu Binarization to separate the background from the tissue. Subsequently, non-overlapping patches of size 256×256 are extracted at a magnification of 20x. These patches are then inputted into a ResNet-18, trained with a contrastive strategy akin to^37^, generating a 1024-dimensional feature vector *h* ∈ ℝ^1024^ for each patch. The set of features 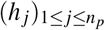 associated with a set *W*_*i*_ comprising *n*_*p*_ patches is consolidated into a feature matrix 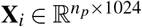. Each patch *x* _*j*_ is defined by its ResNet-18 feature representation *h* _*j*_, encapsulating morphological properties, and a set of coordinates *g* _*j*_ = (*g*_*x, j*_, *g*_*y, j*_) representing the spatial center of the patch. Following the methodology of^38^, a patch filtering process is executed to summarize information in the Whole Slide Image (WSI). This involves Agglomerative Clustering, where a similarity kernel is computed between all patch pairs. Two similarity matrices, 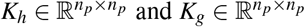 are computed, where 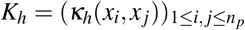 and 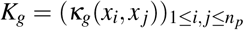. Here, 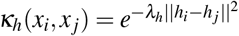 represents a morphological similarity metric, and 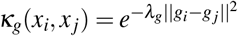 signifies a spatial proximity metric. The overall similarity kernel, *κ*(*x*_*i*_, *x* _*j*_) = *κ*_*h*_(*h*_*i*_, *h* _*j*_)*κ*_*g*_(*g*_*i*_, *g* _*j*_), is employed to group patches with high similarities into clusters denoted as *C*_*k*_. These clusters are then aggregated into a single representation 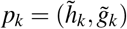, where 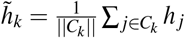 and 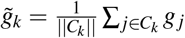. This new patch representation approximates a localized region, effectively reducing the number of instances in Whole Slide Images while preserving a substantial amount of information. We then divide the new patches 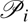 into K categories of regions obtained with a clustering algorithm and interpreted by a pathologist to localize multiple regions of interest in the WSI. This new set of approximated patches will be a stepping stone for constructing a hypergraph denoted by *G*_*i*_ =*< V*_*i*_, *E*_*i*_, **X**_*i*_ *>*. The hyperedges are indicated by an incidence matrix 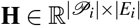 such that,

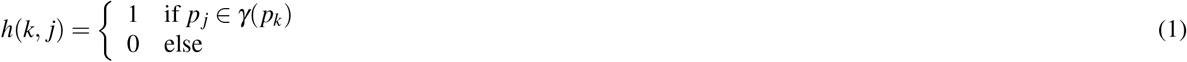

where *γ* (*p*_*j*_) = {*p*_*k*_ ∈ 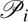; *κ*_*h*_(*p*_*k*_, *p* _*j*_) *≥ δ*_*h*_} and defines the neighborhood around the instance *p* _*j*_. This hypergraph is then fed into a Graph Neural Network composed of a series of hypergraph convolutions and attentions^39^ The resulting feature vector 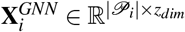 will then be pooled according to each one of the *K* regions so that the final WSI bag becomes 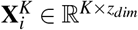.

### 4.2 Integration of Multi-Omics Data

We consider *S* omic sources denoted in the set *O*_*i*_ = *O*_*i,s*_1 *≤ s ≤ S*. The initial step in processing these omic sources involves partitioning each source into P gene sets, each possessing distinct functional properties, defined as *Oi, s* = *O*_*i,s,p*,1*≤p≤P*_. Subsequently, we engage in Variational Autoencoder-based representation learning for each of these P distinct gene sets, a concept introduced in^13^. The encoding networks (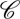 *p*)1 ≤ *p* ≤ *P* are designed to accept inputs from all omic sources about each gene set, encapsulated by the operation 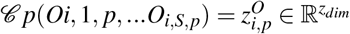. These representations are then concatenated to form a final matrix 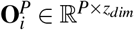, serving as the comprehensive collection of the multi-omics data.

### 4.3 Multimodal Representation and Prediction

Upon acquiring the Whole Slide Image (WSI) bag 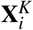 and the multi-omics bag 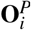, the subsequent phase involves constructing the final multimodal representation. This representation is crafted via a bilinear operation between each pair of elements from both bags, generating a 3-dimensional tensor 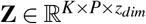 where 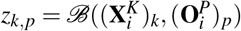 and ℬ signifies the bilinear fusion operator.

We adopt the Multimodal factorized bilinear pooling method introduced in Yu et al.^40^ to execute a meaningful bilinear fusion and encapsulate complex interactions among multimodal features.

Once the final representation materializes, the multimodal tensor is channeled into a downstream network 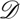 responsible for producing the model’s final prediction. This downstream network comprises a hierarchical mixture of expert networks operating in two distinct stages.

The initial stage involves pathway prediction optimization. For each pathway, *p*, the corresponding spatial regions are inputted into a mixture of experts network^12^. This network yields a pathway representation 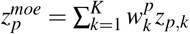, where 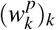 with 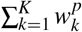 are trainable parameters. Subsequently, these representations are individually directed into single linear layers, each responsible for producing predictions for their respective pathways.

The second stage entails consolidating the representations from all pathways by inputting them into a second Mixture-of-Experts network. This network endeavors to aggregate the pathways into a unified representation 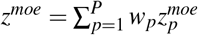, where (*w*_*p*_)*p* with ∑ *p* = 1^*P*^*w*_*p*_ are trainable parameters. The representation is then passed through a single linear layer to generate the final model prediction.

### 4.4 Downstream Tasks

CustOmics accommodates training for three distinct tasks. Firstly, a supervised classification task aims to predict the probability of each class occurrence. This task undergoes training employing a standard Categorical Cross-Entropy loss computed between the predicted classes and the ground truth labels.

The second task involves predicting survival outcomes, trained using the DeepSurv loss function outlined in Katzman et al.^41^. The model adopts the negative partial log-likelihood formula, expressed in our context as:

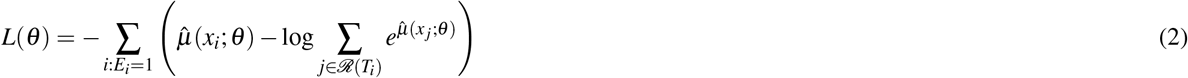

where *E*_*i*_ represents the event for patient *i*, 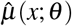 denotes the risk function associated with the risk score estimated by the network’s output layer, and ℛ (*t*) defines the risk set, signifying the patients still susceptible to failure after time *t*.

### 4.5 Multimodal Dropout

To better enforce the robustness of our method to missing data, we implement multimodal dropout introduced in Cheerla et al.^42^ to deal with missing modalities under the assumption of modalities being missing at random. Instead of dropping single neurons, the idea is to drop entire feature vectors corresponding to specific modalities so that it scales up the weights of the others. This is applied to each sample data with a probability *p* for each modality. The dropout rate is a hyperparameter to tune.

### 4.6 Interpretability

To make the results of the CustOmics model interpretable, we implement multiple scores to understand the predictions at different levels of the integration process.

#### 4.6.1 Gene Importance & Pathway Enrichment

In pursuit of enhancing gene-level interpretability, we adapt a method introduced in Withnell et al.^43^ to compute SHAP (Shapley Additive Explanations) values for deep variational autoencoders, as described in^44^. Extending this methodology to the multimodal setting, akin to the approach in Benkirane et al.^13^, becomes paramount.

Post-training the CustOmics network, SHAP values corresponding to genes or latent dimensions are computed for the multi-omics embedding segment. These values estimate the overall contribution by averaging them across samples sharing similar features. The computed SHAP values offer insights at different training phases, delineating gene importance across single-omic, multi-omics, and multimodal integration. Comprehensive elucidation of these processes is available in Text S1.

Furthermore, aiming for a more biologically interpretable model, we propose the derivation of a Pathway Enrichment Score (PES) to assess the impact of specific pathway activations in predictive tasks. Leveraging the weights learned by the gating networks of the mixture of experts, we define a ranking score *r*_*ip*_ per patient and pathway. This score (*r*_*ip*_ = (*w*_*p*_)_*i*_) gauges the overall pathway contribution to the final prediction, forming the basis for pathway ranking for enrichment analysis.

Upon computing these scores, we draw inspiration from Lundberg et al.^45^ to conduct Gene Set Variation Analysis (GSVA) using the importance scores. For each patient *i*, we denote *s*_*i,g*_ as the SHAP value computed by CustOmics. To ensure generalization across all pathways, these values are normalized by the pathway importance computed for the MoE representation, resulting in the normalized SHAP value 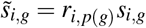, where *p*(*g*) represents the pathway corresponding to the gene *g*.

Subsequently, these normalized scores serve as rankings for computing associated p-values for each pathway. This is achieved by employing a Kolmogorov-Smirnov test, comparing the pathway’s and other genes’ distributions.

#### 4.6.2 Multimodal Interaction Score

To evaluate the importance of the interaction between a spatial region and a functional group, we compute a Multimodal Interaction Score (MIS). This score is directly obtained from the weights of the gating network such that for a specific patient *i*, we have 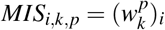.The score measures the impact of a multimodal interaction between a spatial region and a functional group on the final prediction.

### 4.7 Implementation Details

The CustOmics framework is based on the Pytorch deep-learning library^46^. It can be applied to any combination of high-dimensional datasets and histopathology images with multitask training. As done in *Zhang et al*.^26^, DNA methylation data can be divided into 23 separate blocks, each feeding a hidden layer corresponding to a chromosome to avoid overfitting and save GPU memory.

Inspired by the work in^13^, we adopt a multiphase training strategy to ensure the optimal integration of all modalities. During the first phase, we train each modality independently to obtain unsupervised sub-representations for the different bags. The second phase consists of unfreezing the central encoding network that learns the supervised representation of the final bag.

The whole architecture is built using fully connected blocks with weights initialized following a uniform distribution 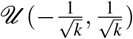 where *k* is the number of weight parameters. We use a batch normalization technique in each layer composing the neural network to address the internal covariate shift problem^47^. Also, to avoid overfitting problems, we use dropout^48^; its rate is considered a hyperparameter.

The input dataset was randomly split into training, validation, and testing sets (60-20-20%) using stratified 5-fold cross-validation so that the proportion of samples in each tumor type between the different sets is preserved in all the folds. We perform Bayesian optimization^49^ using the validation set to find our model’s best possible combination of hyperparameters.

## Supplementary information

### 4.8 Supplementary Tables

### 4.9 Supplementary Texts

#### Text S1: SHAP Values

SHAP (Shapley Additive exPlanations) values provide a method to explain the output of any machine learning model grounded in cooperative game theory. The concept is based on Shapley values, which distribute ‘payouts’ fairly to ‘players’ in a coalition. In machine learning, ‘payouts’ are the model’s predictions, and ‘players’ are the input features.

The Shapley value, a core concept in cooperative game theory, offers a fair distribution of payoffs among players based on their contribution. For a player *i* in a set *N* of players, the Shapley value is mathematically defined as:

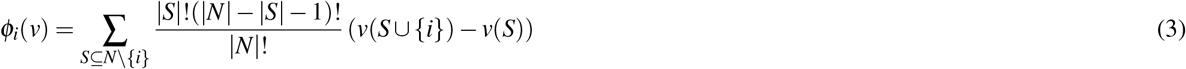

**Table 3.** Table S1: Datasets Description. Description of the TCGA datasets used in this study for both classification and survival tasks. We show the number of patients available for each modality along with the censoring rate of the cohort.

|  |  | Missing | Mutations | CNV | RNAseq | Methylation | WSI | Overall |
| --- | --- | --- | --- | --- | --- | --- | --- | --- |
| n |  |  |  |  |  |  |  | 19188 |
| program, n (%) | TARGET | 0 |  |  |  |  |  | 4666 (24.3) |
|  | TCGA |  |  |  |  |  |  | 14522 (75.7) |
| sample_type, n (%) | Additional - New Primary | 71 |  |  |  |  |  | 11 (0.1) |
|  | Additional Metastatic |  |  |  |  |  |  | 2 (0.0) |
|  | Blood Derived Normal |  |  |  |  |  |  | 2 (0.0) |
|  | Bone Marrow Normal |  |  |  |  |  |  | 845 (4.4) |
|  | Buccal Cell Normal |  |  |  |  |  |  | 5 (0.0) |
|  | FFPE Scrolls |  |  |  |  |  |  | 10 (0.1) |
|  | Human Tumor Original Cells |  |  |  |  |  |  | 2 (0.0) |
|  | Metastatic |  |  |  |  |  |  | 411 (2.1) |
|  | Primary Blood Derived Cancer - Bone Marrow |  |  |  |  |  |  | 1054 (5.5) |
|  | Primary Blood Derived Cancer - Peripheral Blood |  |  |  |  |  |  | 616 (3.2) |
|  | Primary Tumor |  |  |  |  |  |  | 13060 (68.3) |
|  | Recurrent Blood Derived Cancer - Bone Marrow |  |  |  |  |  |  | 208 (1.1) |
|  | Recurrent Blood Derived Cancer - Peripheral Blood |  |  |  |  |  |  | 14 (0.1) |
|  | Recurrent Tumor |  |  |  |  |  |  | 103 (0.5) |
|  | Solid Tissue Normal |  |  |  |  |  |  | 2774 (14.5) |
| project_id, n (%) | TARGET-ALL-P3 | 234 | 0 | 0 | 0 | 0 | 0 | 341 (1.8) |
|  | TARGET-AML |  | 0 | 0 | 0 | 0 | 0 | 2029 (10.7) |
|  | TARGET-CCSK |  | 0 | 0 | 0 | 0 | 0 | 19 (0.1) |
|  | TARGET-NBL |  | 0 | 0 | 0 | 0 | 0 | 1109 (5.9) |
|  | TARGET-OS |  | 0 | 0 | 0 | 0 | 0 | 288 (1.5) |
|  | TARGET-RT |  | 0 | 0 | 0 | 0 | 0 | 61 (0.3) |
|  | TARGET-WT |  | 0 | 0 | 0 | 0 | 0 | 789 (4.2) |
|  | TCGA-ACC |  | 89 | 89 | 80 | 80 | 227 | 97 (0.5) |
|  | TCGA-BLCA |  | 408 | 408 | 426 | 431 | 457 | 454 (2.4) |
|  | TCGA-BRCA |  | 1082 | 1082 | 1179 | 881 | 1133 | 1283 (6.8) |
|  | TCGA-CESC |  | 284 | 284 | 299 | 299 | 279 | 317 (1.7) |
|  | TCGA-CHOL |  | 36 | 36 | 45 | 45 | 39 | 71 (0.4) |
|  | TCGA-COAD |  | 442 | 442 | 436 | 335 | 459 | 571 (3.0) |
|  | TCGA-DLBC |  | 47 | 47 | 46 | 47 | 44 | 52 (0.3) |
|  | TCGA-ESCA |  | 184 | 184 | 197 | 201 | 158 | 251 (1.3) |
|  | TCGA-GBM |  | 606 | 606 |  | 151 | 860 | 671 (3.5) |
|  | TCGA-HNSC |  | 523 | 523 | 568 | 579 | 472 | 612 (3.2) |
|  | TCGA-KICH |  | 65 | 65 | 89 | 65 | 121 | 184 (1.0) |
|  | TCGA-KIRC |  | 532 | 532 | 587 | 478 | 519 | 985 (5.2) |
|  | TCGA-KIRP |  | 284 | 284 | 321 | 316 | 300 | 381 (2.0) |
|  | TCGA-LAML |  | 170 | 170 | 164 | 120 |  | 697 (3.7) |
|  | TCGA-LGG |  | 528 | 528 | 525 | 529 | 844 | 538 (2.8) |
|  | TCGA-LIHC |  | 372 | 372 | 419 | 424 | 379 | 469 (2.5) |
|  | TCGA-LUAD |  | 508 | 508 | 541 | 482 | 541 | 877 (4.6) |
|  | TCGA-LUSC |  | 490 | 490 | 511 | 399 | 512 | 765 (4.0) |
|  | TCGA-MESO |  | 85 | 85 | 85 | 85 | 87 | 88 (0.5) |
|  | TCGA-OV |  | 587 | 587 | 494 | 10 | 107 | 758 (4.0) |
|  | TCGA-PAAD |  | 184 | 184 | 182 | 194 | 209 | 223 (1.2) |
|  | TCGA-PCPG |  | 169 | 169 | 187 | 187 | 196 | 189 (1.0) |
|  | TCGA-PRAD |  | 502 | 502 | 551 | 553 | 449 | 623 (3.3) |
|  | TCGA-READ |  | 157 | 157 | 156 | 98 | 166 | 192 (1.0) |
|  | TCGA-SARC |  | 261 | 261 | 260 | 266 | 600 | 290 (1.5) |
|  | TCGA-SKCM |  | 458 | 458 | 439 | 461 | 475 | 477 (2.5) |
|  | TCGA-STAD |  | 408 | 408 | 444 | 382 | 442 | 544 (2.9) |
|  | TCGA-TGCT |  | 139 | 139 | 139 | 139 | 254 | 156 (0.8) |
|  | TCGA-THCA |  | 511 | 511 | 572 | 570 | 519 | 615 (3.2) |
|  | TCGA-THYM |  | 123 | 123 | 125 | 125 | 181 | 139 (0.7) |
|  | TCGA-UCEC |  | 538 | 538 | 558 | 465 | 566 | 606 (3.2) |
|  | TCGA-UCS |  | 54 | 54 | 55 | 55 | 91 | 63 (0.3) |
|  | TCGA-UVM |  | 80 | 80 | 80 | 80 | 80 | 80 (0.4) |
| Age at Diagnosis in Years, mean (SD) |  | 511 |  |  |  |  |  | 46.5 (26.0) |
| Gender, n (%) | Female | 450 |  |  |  |  |  | 34 (0.2) |
|  | Male |  |  |  |  |  |  | 27 (0.1) |
|  | Female |  |  |  |  |  |  | 9342 (49.9) |
|  | Female |  |  |  |  |  |  | 6 (0.0) |
|  | Male |  |  |  |  |  |  | 9311 (49.7) |
|  | Male |  |  |  |  |  |  | 12 (0.1) |
|  | Unknown |  |  |  |  |  |  | 1 (0.0) |
|  | not reported |  |  |  |  |  |  | 5 (0.0) |

where *S* is a subset of players excluding *i*, and *v*(*S*) is the value function of the coalition *S*.

In machine learning, SHAP values interpret the contribution of each feature to predicting a particular instance. The SHAP value of feature *j* for a prediction instance *x* in a feature set *X* is calculated as:

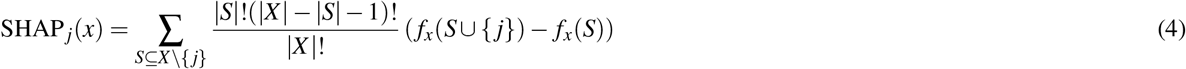

where *f*_*x*_(*S*) represents the output of model *f* when the feature set *S* is present for instance *x*.

SHAP values offer several advantages in model interpretation:

**Table 4.**
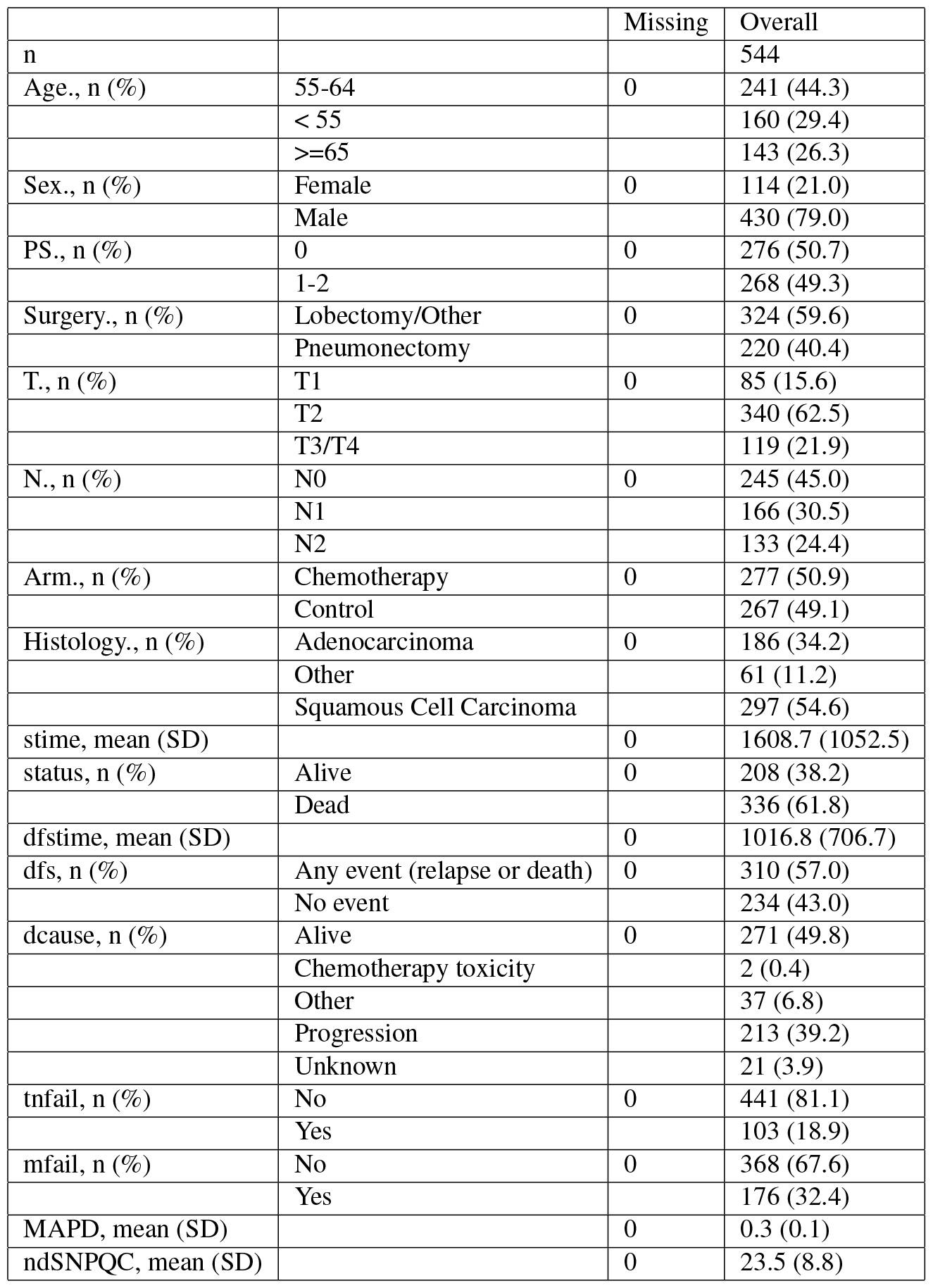
Table S2: IALT Overview. Overview of the IALT cohort with details about the different clinical and molecular features involved.

#### Consistency

If the model changes such that a feature’s contribution increases or remains the same, its SHAP value should not decrease.

#### Local Accuracy

The sum of SHAP values for all features plus a base value equals the model prediction.

#### Global Interpretability

Aggregating SHAP values across a dataset provides an overview of the model’s behavior.

Applicable to any machine learning model, SHAP values are precious in areas requiring high transparency, such as healthcare, finance, and criminal justice. They enhance the understanding and trustworthiness of AI-driven decisions by offering detailed insights into the contributions of individual features.

#### Text S2: Survival Analysis

Survival analysis is a branch of statistics focused on analyzing time until an event of interest, commonly used in diverse fields like medicine, biology, and engineering. This analysis deals with the challenge of censored data, where the event time for some subjects is unknown.

At the core of survival analysis is the **Survival Function** *S*(*t*), representing the probability of survival beyond time *t*. Defined as *S*(*t*) = *P*(*T > t*), where *T* represents the survival time, this function is characterized by its non-increasing nature, starting from *S*(0) = 1 and approaching zero as time goes to infinity.

**Table 5.** Table S3: Modality Combinations. Performance comparison between multiple combination of modalities for CustOmics for the Pancancer classification task. The evaluation is done using the Area Under ROC-curve (AUC).

| Omics Combinations | SNN | CustOmics |
| --- | --- | --- |
| CNV | 74.3 ± 3.0 | 75.1 ± 2.7 |
| RNAseq | 94.0 ± 2.6 | 96.0 ± 1.4 |
| Methyl | 81.1 ± 1.7 | 82.3 ± 1.3 |
| CNV + RNAseq | 94.3 ± 2.9 | 96.9 ± 0.8 |
| CNV + Methyl | 81.4 ± 1.8 | 85.7 ± 2.1 |
| RNAseq + methyl | 93.2 ± 1.0 | 97.3 ± 0.7 |
| CNV + RNAseq + Methyl | 94.1 ± 2.7 | 98.9 ± 1.4 |

**Table 6.** Table S4: Ablation Study. Performance comparison between CustOmics and the state of the art for classification tasks by replacing different instances of the model: **a. Multi-Omics Ablation:** CustOmics A1 replaces the multi-omics VAE with an SNN. **b. Hypergraph Ablation:** CustOmics A2 replaces the hypergraph encoder with a visual transformer and CustOmics A3 replaces the hypergraph represnetation with a regular graph embedding. **c. Downstream Network Ablation:** CustOmics A4 replaces the hierarchical mixture-of-experts approach with a regular mixture of experts network, CustOmics A5 replaces it with a transformer classifier.

| Methods | PANCAN | BRCA | COAD | STAD |
| --- | --- | --- | --- | --- |
| SNN (Multi-Omics) | 94.1 ± 2.7 | 92.0 ± 3.3 | 79.2 ± 3.3 | 84.6 ± 3.3 |
| CustOmics (Multi-Omics) | **98.9 ± 1.4** | **98.3 ± 1.0** | **88.1 ± 1.9** | **98.4 ± 1.2** |
| DeepSets (WSI Only) | 84.7 ± 3.3 | 68.6 ± 4.1 | 55.2 ± 4.3 | 58.9 ± 4.6 |
| AttnMIL (WSI Only) | 88.4 ± 2.0 | 71.2 ± 4.9 | 56.2 ± 4.4 | 61.4 ± 4.4 |
| DeepAttnMISL (WSI Only) | 89.8 ± 2.5 | 71.1 ± 3.3 | 55.7 ± 4.0 | 62.1 ± 4.5 |
| MCAT (WSI Only) | 90.4 ± 1.8 | 72.3 ± 3.3 | 61.7 ± 3.3 | 69.4 ± 3.5 |
| CustOmics A1 (WSI Only) | 85.1 ± 3.8 | 70.5 ± 5.5 | 57.7 ± 4.1 | 59.4 ± 4.4 |
| CustOmics A2 (WSI Only) | 90.4 ± 1.8 | 72.3 ± 3.3 | 61.7 ± 3.3 | 69.4 ± 3.5 |
| CustOmics A3 (WSI Only) | 88.7 ± 1.2 | 69.4 ± 3.6 | 55.1 ± 3.5 | 61.9 ± 5.0 |
| CustOmics A4 (WSI Only) | 89.4 ± 2.1 | 71.0 ± 3.3 | 58.8 ± 3.3 | 67.2 ± 3.7 |
| CustOmics A5 (WSI Only) | 87.1 ± 1.9 | 70.4 ± 3.6 | 55.2 ± 3.9 | 66.4 ± 4.1 |
| DeepSets (WSI + Multi-Omics) | 96.7 ± 1.5 | 84.9 ± 2.0 | 58.6 ± 2.2 | 67.1 ± 2.7 |
| AttnMIL (WSI + Multi-Omics) | 97.1 ± 1.2 | 86.6 ± 2.1 | 60.0 ± 2.5 | 69.4 ± 2.7 |
| DeepAttnMISL (WSI + Multi-Omics) | 97.8 ± 1.1 | 88.3 ± 2.7 | 65.4 ± 2.7 | 66.6 ± 2.2 |
| MCAT (WSI + Multi-Omics) | 98.9 ± 1.1 | 95.4 ± 2.0 | 93.3 ± 1.2 | 88.7 ± 2.3 |
| CustOmics A1 (WSI + Multi-Omics) | 96.1 ± 1.7 | 85.2 ± 2.4 | 59.5 ± 1.9 | 67.7 ± 2.9 |
| CustOmics A2 (WSI + Multi-Omics) | 99.0 ± 1.3 | 98.4 ± 2.2 | 94.1 ± 1.2 | 95.3 ± 2.4 |
| CustOmics A3 (WSI + Multi-Omics) | 99.9 ± 1.1 | 98.2 ± 2.5 | 93.9 ± 1.9 | 94.1 ± 2.7 |
| CustOmics A4 (WSI + Multi-Omics) | 99.4 ± 0.8 | 98.0 ± 1.2 | 94.2 ± 1.1 | 96.1 ± 2.2 |
| CustOmics A5 (WSI + Multi-Omics) | 98.5 ± 1.9 | 97.3 ± 3.4 | 93.8 ± 2.0 | 94.2 ± 2.9 |

Complementing the survival function is the **Hazard Function** λ (*t*), which provides the instantaneous event rate at time *t*, conditional on survival until that point. It is mathematically expressed as:

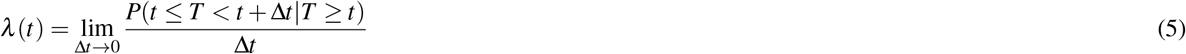

This function plays a pivotal role in understanding the dynamics of the time-to-event process.

Handling incomplete data due to censoring is a crucial aspect of survival analysis. Censoring occurs when the information about an individual’s event time is incomplete, and it comes in various forms, such as right-censoring, left-censoring, and interval-censoring.

The **Kaplan-Meier Estimator** is a cornerstone in non-parametric survival analysis, used for estimating the survival function from life-table data. Given observed survival times, this estimator calculates the survival probability at different time points, accounting for censored data.

For examining the relationship between survival time and one or more predictor variables, the **Cox Proportional Hazards Model** is extensively used. It is a semi-parametric model that defines the hazard function as:

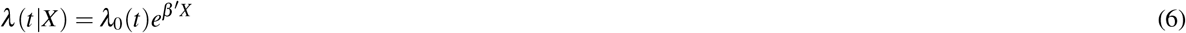

where λ _0_(*t*) is the baseline hazard, *X* represents covariates, and *β* is the coefficient vector.

In cases where a specific distribution of survival times is assumed, **Parametric Models** like exponential, Weibull or log-normal models are employed. These models offer greater flexibility but require conforming to certain distributional assumptions.

Survival analysis is indispensable across various applications, from assessing patient survival in medical studies to analyzing failure times in engineering. Its capacity to handle censored data and model time-to-event relationships makes it a vital statistical analysis and research tool.

In summary, survival analysis provides a comprehensive framework for analyzing and predicting the time until an event of interest occurs. Using different statistical methods, it addresses the complexities of censored data and offers insights into the factors influencing survival times.

#### Text S3: Mixture of Experts

The Mixture of Experts (MoE) model is a versatile approach in statistical modeling and machine learning, designed to capture complex patterns in data by combining the predictions of multiple ‘expert’ models. This approach is particularly effective in scenarios where different subsets of data are best explained by different types of models.

Central to the MoE model is the idea of partitioning the input space into regions, each dominated by an expert, which is typically a separate model or learner. The combination of these experts’ predictions provides a comprehensive output that reflects the nuances of the entire dataset.

The architecture of a MoE model comprises two main components:

#### Experts

Each expert is a model (like a regression or a neural network) trained on the dataset. In a MoE model, multiple experts are trained to specialize in different parts of the input space.

#### Gating Network

This is a probabilistic model that determines the weight or influence of each expert for a given input. The gating network’s output is a set of weights that sum to one, indicating the proportion of contribution from each expert.

Mathematically, the output of a MoE model for an input *x* is a weighted sum of the outputs from the experts. If *y*_*i*_(*x*) is the output of the *i*-th expert and *g*_*i*_(*x*) is the weight assigned by the gating network for this expert, the overall output *Y* (*x*) is given by:

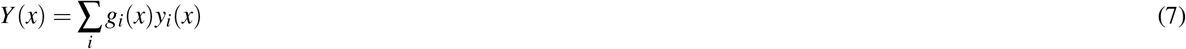

where ∑_*i*_ *g*_*i*_(*x*) = 1, and *g*_*i*_(*x*) is typically determined using a softmax function over the gating network outputs.

One of the key advantages of the MoE model is its flexibility. By combining experts that are proficient in different regions of the input space, the MoE model can approximate a wide variety of functions, making it suitable for complex tasks like pattern recognition, time-series forecasting, and nonlinear regression.

In practical applications, the training of a MoE model involves optimizing both the parameters of the experts and the gating network, typically using gradient-based methods. This process involves balancing the specialization of each expert with the overall coherence of the model.

## References

1. Hosny, A., Parmar, C., Quackenbush, J., Schwartz, L. H. & Aerts, H. J. Artificial intelligence in radiology. Nat. Rev. Cancer 18, 500–510 (2018).

2. Vamathevan, J. et al. Applications of machine learning in drug discovery and development. Nat. reviews Drug discovery 18, 463–477 (2019).

3. Bera, K., Schalper, K. A., Rimm, D. L., Velcheti, V. & Madabhushi, A. Artificial intelligence in digital pathology—new tools for diagnosis and precision oncology. Nat. reviews Clin. oncology 16, 703–715 (2019).

4. Zitnik, M. et al. Machine learning for integrating data in biology and medicine: Principles, practice, and opportunities. Inf. Fusion 50, 71–91 (2019).

5. Ho, W. et al. Multi-omic profiling of lung and liver tumor microenvironments of metastatic pancreatic cancer reveals site-specific immune regulatory pathways. Genome Biol. 22, DOI: 10.1186/s13059-021-02363-6 (2021).

6. Hira, M. et al. Integrated multi-omics analysis of ovarian cancer using variational autoencoders. Sci. Reports 11, 6265 (2021).

7. Suzuki, K. Overview of deep learning in medical imaging. Radiol. physics technology 10, 257–273 (2017).

8. Kim, M. et al. Deep learning in medical imaging. Neurospine 16, 657 (2019).

9. Hanna, M. G., Parwani, A. & Sirintrapun, S. J. Whole slide imaging: technology and applications. Adv. Anat. Pathol. 27, 251–259 (2020).

10. Sudharshan, P. et al. Multiple instance learning for histopathological breast cancer image classification. Expert. Syst. with Appl. 117, 103–111 (2019).

11. Shao, Z. et al. Transmil: Transformer based correlated multiple instance learning for whole slide image classification. Adv. Neural Inf. Process. Syst. 34, 2136–2147 (2021).

12. Jordan, M. I. & Jacobs, R. A. Hierarchical mixtures of experts and the em algorithm. Neural computation 6, 181–214 (1994).

13. Benkirane, H., Pradat, Y., Michiels, S. & Cournède, P.-H. Customics: A versatile deep-learning based strategy for multi-omics integration. PLoS Comput. Biol. 19, e1010921 (2023).

14. Chen, R. J. et al. Multimodal co-attention transformer for survival prediction in gigapixel whole slide images. In Proceedings of the IEEE/CVF International Conference on Computer Vision, 4015–4025 (2021).

15. Klambauer, G., Unterthiner, T., Mayr, A. & Hochreiter, S. Self-normalizing neural networks. Adv. neural information processing systems 30 (2017).

16. Chen, R. J. et al. Pan-cancer integrative histology-genomic analysis via multimodal deep learning. Cancer Cell 40, 865–878 (2022).

17. Zaheer, M. et al. Deep sets. Adv. neural information processing systems 30 (2017).

18. Ilse, M., Tomczak, J. & Welling, M. Attention-based deep multiple instance learning. In International conference on machine learning, 2127–2136 (PMLR, 2018).

19. Yao, J., Zhu, X., Jonnagaddala, J., Hawkins, N. & Huang, J. Whole slide images based cancer survival prediction using attention guided deep multiple instance learning networks. Med. Image Analysis 65, 101789 (2020).

20. Graham, S. et al. Hover-net: Simultaneous segmentation and classification of nuclei in multi-tissue histology images. Med. image analysis 58, 101563 (2019).

21. Saltz, J. et al. Spatial organization and molecular correlation of tumor-infiltrating lymphocytes using deep learning on pathology images. Cell reports 23, 181–193 (2018).

22. Grossman, R. L. et al. Toward a shared vision for cancer genomic data. New Engl. J. Medicine 375, 1109–12 (2016).

23. Group, I. A. L. C. T. C. Cisplatin-based adjuvant chemotherapy in patients with completely resected non–small-cell lung cancer. New Engl. J. Medicine 350, 351–360 (2004).

24. Hand, D. J. & Till, R. J. A simple generalisation of the area under the roc curve for multiple class classification problems. Mach. learning 45, 171–186 (2001).

25. Zhang, X. et al. Integrated multi-omics analysis using variational autoencoders: Application to pan-cancer classification. In 2019 IEEE International Conference on Bioinformatics and Biomedicine (BIBM), 765–69 (2019).

26. Zhang, X., Xing, Y., Sun, K. & Guo, Y. Omiembed: A unified multi-task deep learning framework for multi-omics data. Cancers 13, 3047 (2021).

27. Mehta, A. & Tripathy, D. Co-targeting estrogen receptor and her2 pathways in breast cancer. The breast 23, 2–9 (2014).

28. Cruz, R. G., Madden, S. F., Brennan, K. & Hopkins, A. M. A transcriptional link between her2, jam-a and foxa1 in breast cancer. Cells 11, 735 (2022).

29. Tape, C. J. et al. Oncogenic kras regulates tumor cell signaling via stromal reciprocation. Cell 165, 910–920 (2016).

30. El Bairi, K. et al. The tale of tils in breast cancer: a report from the international immuno-oncology biomarker working group. NPJ Breast Cancer 7, 150 (2021).

31. Loi, S. et al. The journey of tumor-infiltrating lymphocytes as a biomarker in breast cancer: clinical utility in an era of checkpoint inhibition. Annals Oncol. 32, 1236–1244 (2021).

32. Zhao, H. et al. Inflammation and tumor progression: signaling pathways and targeted intervention. Signal transduction targeted therapy 6, 263 (2021).

33. Mahdizadehi, M., Saghaeian Jazi, M., Mir, S. M. & Jafari, S. M. Role of fibrilins in human cancer: A narrative review. Heal. Sci. Reports 6, e1434 (2023).

34. Sunaga, N. et al. Knockdown of oncogenic kras in non–small cell lung cancers suppresses tumor growth and sensitizes tumor cells to targeted therapy. Mol. cancer therapeutics 10, 336–346 (2011).

35. Liu, C. et al. The superior efficacy of anti-pd-1/pd-l1 immunotherapy in kras-mutant non-small cell lung cancer that correlates with an inflammatory phenotype and increased immunogenicity. Cancer letters 470, 95–105 (2020).

36. Lu, M. Y. et al. Data-efficient and weakly supervised computational pathology on whole-slide images. Nat. biomedical engineering 5, 555–570 (2021).

37. Ciga, O., Xu, T. & Martel, A. L. Self supervised contrastive learning for digital histopathology. Mach. Learn. with Appl. 7, 100198 (2022).

38. Benkirane, H. et al. Hyper-adac: Adaptive clustering-based hypergraph representation of whole slide images for survival analysis. In Parziale, A. et al. (eds.) Proceedings of the 2nd Machine Learning for Health symposium, vol. 193 of Proceedings of Machine Learning Research, 405–418 (PMLR, 2022).

39. Bai, S., Zhang, F. & Torr, P. H. Hypergraph convolution and hypergraph attention. Pattern Recognit. 110, 107637 (2021).

40. Yu, Z., Yu, J., Xiang, C., Fan, J. & Tao, D. Beyond bilinear: Generalized multimodal factorized high-order pooling for visual question answering. IEEE Transactions on Neural Networks Learn. Syst. 29, 5947–5959 (2018).

41. Katzman, J. et al. Deepsurv: Personalized treatment recommender system using a cox proportional hazards deep neural network. BMC Med. Res. Methodol. 18, 24 (2018).

42. Cheerla, A. & Gevaert, O. Deep learning with multimodal representation for pancancer prognosis prediction. Bioinformatics 35, i446–i454 (2019).

43. Withnell, E., Zhang, X., Sun, K. & Guo, Y. Xomivae: an interpretable deep learning model for cancer classification using high-dimensional omics data. Briefings Bioinforma. 22, 315 (2021).

44. Lundberg, S. M. & Lee, S.-I. A unified approach to interpreting model predictions. In Guyon, I.et al. (eds.) Advances in Neural Information Processing Systems 30, 4765–4774 (Curran Associates, Inc., 2017).

45. Hänzelmann, S., Castelo, R. & Guinney, J. Gsva: gene set variation analysis for microarray and rna-seq data. BMC bioinformatics 14, 1–15 (2013).

46. Paszke, A. et al. Pytorch: An imperative style, high-performance deep learning library. In Advances in Neural Information Processing Systems 32, 8024–8035 (Curran Associates, Inc., 2019).

47. Ioffe, S. & Szegedy, C. Batch normalization: Accelerating deep network training by reducing internal covariate shift. In International conference on machine learning, 448–456 (PMLR, 2015).

48. Srivastava, N., Hinton, G., Krizhevsky, A., Sutskever, I. & Salakhutdinov, R. Dropout: A simple way to prevent neural networks from overfitting. J. Mach. Learn. Res. 15, 1929–1958 (2014).

49. Snoek, J., Larochelle, H. & Adams, R. P. Practical bayesian optimization of machine learning algorithms. In Proceedings of the 25th International Conference on Neural Information Processing Systems - Volume 2 (2012).

